# Estimating amino acid substitution models from genome datasets: A simulation study on the performance of estimated models

**DOI:** 10.1101/2023.04.09.536188

**Authors:** Tinh Nguyen Huy, Cuong Cao Dang, Le Sy Vinh

## Abstract

Estimating amino acid substitution models is a crucial task in bioinformatics. The maximum likelihood (ML) approach has been proposed to estimate amino acid substitution models from large datasets. The quality of newly estimated models is normally assessed by comparing with the existing models in building ML trees. Two important questions remained are the correlation of the estimated models with the true models and the required size of the training datasets to estimate reliable models. In this paper, we performed a simulation study to answer these two questions based on the simulated data. We simulated genome datasets with different number of genes/alignments based on predefined models (called true models) and predefined trees (called true trees). The simulated datasets were used to estimate amino acid substitution model using the ML estimation method. Our experiment showed that models estimated by the ML methods from simulated datasets with more than 100 genes have high correlations with the true models. The estimated models performed well in building ML trees in comparison with the true models. The results suggest that amino acid substitution models estimated by the ML methods from large genome datasets might play as reliable tool for analyzing amino acid sequences.

## Introduction

The amino acid (AA) substitution model describes the substitution rates among twenty amino acids during the evolution. The substitutions among amino acids are typically modeled by a time-continuous, time-homologous, and stationary Markov process. The model is continuous in time indicating that the AA substitutions can happen at any point during the evolution process. The time-homologous property assumes that the substitution rates among amino acids remains constant through the entire process. The stationary property requires the stationary of amino acid frequencies during the evolution. The AA substitution process might be assumed to be time-reversible indicating that the exchangeability rates between two amino acids are the same in both directions. This assumption reduces the computational effort in estimating and using the models, however, the time-reversible property might be not biologically realistic and does not allow us to reconstruct rooted trees.

The AA substitution models play a vital role in analyzing the evolutionary relationships among protein sequences. They are used to calculate the likelihood values of phylogenetic trees to determine the maximum likelihood tree. They can be also employed as the score matrices to determine the similarity between protein sequences in sequence similarity searches or multiple sequence alignment algorithms.

The amino acid substitution model contains a large number of parameters, i.e., 189 parameters for time reversible models or 379 parameters for the time non-reversible models. Several maximum likelihood methods have been proposed to estimated AA substitution models (C. C. Dang et al. 2022; Minh et al. 2021; C. C. a. Dang et al. 2014). A big dataset of alignments (called the training dataset) is required to estimate AA substitution models to overcome the overfitting problem when estimating a large number of parameters (Le and Gascuel 2008; Whelan and Goldman 2001; Jones, Taylor, and Thornton 1992; Minh et al. 2021; Vinh 2021).

The time reversible models have been widely used in analyzing protein sequences. The ML estimation method QMaker(Minh et al. 2021) has been introduced to estimate the time-reversible models from whole genome datasets. They applied QMaker to estimate several time reversible models from genomes of plants, birds, yeasts, mammals and insects. The clade-specific models outperform the other existing models in building ML trees for their corresponding data.

The time reversible assumption is mathematically and computationally convenient, however, studies revealed that the assumption may be violated (Squartini and Arndt 2008; Naser-Khdour et al. 2019). Recently, the nQMaker method (C. C. Dang et al. 2022) has been proposed to estimate time non-reversible models from genome datasets. Experiments show that the time non-reversible models are likely better than the time reversible models in analyzing large alignments. Crucially, the time non-reversible models allow us to reconstruct rooted phylogenetic trees.

The quality of estimated models is compared with the existing models in building ML trees (i.e., model M1 is considered better than model M2 if the ML tree constructed with M1 is better than that constructed with M2). For training datasets containing real alignments, the true models and true trees are unknown, therefore, we do not know how the estimated models are closely related with the true models; and if the trees constructed with the estimated models are as good as that constructed with the true models. The size of training datasets to estimate reliable models is also an important open question.

In this paper, we used the alignment simulation program AliSim (Ly-Trong et al. 2022) to simulate various datasets each containing *N* alignments/genes (*N* = 1,2,5,10,20,50,100,200,250) generated from four predefined AA substitution models (Q.plant, NQ.plant, Q.bird, and NQ.bird) and two predefined phylogenetic trees (i.e., the plant tree (Ran et al. 2018) and the bird tree (Jarvis et al. 2015)). The predefined models and phylogenetics are considered/called the true models and true phylogenetic trees of the simulated datasets. For each simulated dataset, we used the QMaker and nQMaker programs (Minh et al. 2021; C. C. Dang et al. 2022) to estimate a time-reversible model and a time non-reversible model, respectively. We performed different analyses to directly compare the estimated models with the true models. We also assessed the ability of the estimated models in building ML trees in comparison with the true models.

## Materials and methods

### Data

We used the AliSim program (Ly-Trong et al. 2022) to simulate amino acid alignments. Typically, AliSim requires five parameters to simulate an alignment: a predefined AA substitution model, a site rate heterogeneous model, a phylogenetic tree, the number of sequences together with the sequence length of the alignment. We simulated alignments with both time reversible substitution models (called time reversible alignments) and time non-reversible substitution models (called time non-reversible alignments).

We used real parameters obtained from the plant genome dataset of 1000 genes, each containing 35 sequences (Ran et al. 2018) and the bird genome dataset of 6295 genes, each consisting of 48 sequences (Jarvis et al. 2015) to create simulated plant and bird genome datasets.

- Simulating plant genome datasets: To create a simulated dataset of *N* alignments/genes (*N* = 1,2,5,10,20,50,100,200,250), we randomly selected *N* genes from the plant genome. For each selected gene, the sequence length, the time reversible substitution model Q.plant, the site rate heterogeneity model (estimated from the selected gene), and an unrooted phylogenetic tree (Minh et al. 2021) were provided to simulate a time reversible alignment. Similarly, we used the time non-reversible substitution model NQ.plant and the rooted phylogenetic tree (C. C. Dang et al. 2022) to simulate a time non-reversible alignment.
- Simulating bird genome datasets: We used the same above procedure to create simulated bird genome datasets each containing *N* alignments (*N* = 1,2,5,10,20,50,100,200,250).

To compare the performance of estimated models and the true models in building ML trees, we used the testing datasets including all 308 plant testing genes and a subset of 500 bird testing genes obtained from the QMaker’s paper (Minh et al. 2021).

## Methods

### Estimating amino acid substitution model

We assume that the evolutionary process is independent among amino acid sites. We use a Markov process to model the AA substitution process at a site with time-homologous, time-continuous, and stationary assumptions. The core component of the model is the so-called *Q* = {*q_xy_*} matrix of 20*x*20 instantaneous substitution rates among amino acids. Technically, *q*_xy_ represents the number of substitutions from amino acid *x* to amino acid *y* per time unit. We normally normalize the matrix *Q* so that the branch length of phylogenetic trees reflects the number of AA substitution per site, i.e., *Q* contains 379 free parameters.

To reduce the computational burdens, we might assume that the amino acid substitution process is time-reversible (i.e., the exchangeability rates between amino acid *x* and amino acid *y* are equal in both directions). As a result, the matrix *Q* can be decomposed into two parts: a symmetric exchangeability rate matrix *R* = {*r_xy_*} and an amino acid frequency vector *⊓* = {*π_x_*} such that *q_xy_* = *π_y_r_xy_* and *q_xx_* = −∑*_y_ q_xy_*. The *R* matrix contains 189 free parameters and the *⊓* vector consists of 19 free parameters. Note that we are unable to root the ML phylogenetic trees with the time reversible models.

The training dataset used to estimate an AA substitution model includes *N* alignments denoted by *D* = {*D*_1_…, *D_N_*}. Let *T* = {*T*_1_…,*T_N_*} be the set of *N* trees for the training alignments, i.e., *T_i_* is the phylogenetic tree of alignment *D_i_*. To account the site rate heterogeneity among sites in the model estimation process, let *P* = {*ρ*_1_,…, *ρ_N_*} be the set of site rate models for the training alignments, i.e., *ρ_i_* is the site rate model for alignment *D_i_*.

The maximum likelihood estimation method determines the parameters of model *Q*, the phylogenetic trees **T**, and parameters of site rate models **P** to maximize the likelihood value *L*(*Q*|*T,P,D*). We can calculate the likelihood value over *N* alignments of **D** as following:

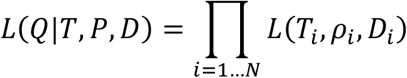

Optimizing *L*(*Q*|*T,P,D*) with a large number of free parameters is a computationally expensive task. Fortunately, the previous studies (Minh et al. 2021; C. C. Dang et al. 2022; Le and Gascuel 2008; C. C. a. Dang et al. 2014) show that the parameters of *Q* can be optimized based on nearly optimal trees **T** and site rate models **P**. Thus, the parameters of *Q*, **T**, and **P** can be estimated iteratively instead of simultaneously to overcome the computational burdens. In this paper, we used the recently published ML estimation method QMaker and nQMaker (Minh et al. 2021; C. C. Dang et al. 2022) to estimate time reversible models and time non-reversible models from genome datasets, respectively.

### Alignment simulation

We used the AliSim program (Ly-Trong et al. 2022) to simulate alignments based on parameters of real alignments. Given a real alignment *R_i_*, the procedure to simulate an alignment *D_i_* ∈ *D* includes two steps (see Figure *1*) described as following:

- Parameter inference step: Consider a real alignment *R_i_*, including *m_i_* sequences with sequence length *l_i_*. If *R_i_*, is selected from plant (bird) genome dataset, assign the predefined substitution model *Q^defined^* by the Q.plant (Q.bird) model, and the predefined tree *T^defined^* by the plant (bird) tree to simulate the alignment *D_i_*. We used the IQ-TREE2 program (Minh et al. 2020) to determine the best site-rate heterogeneity model *ρ_i_* for *R_i_*.
- Simulation step: The AA substitution model *Q^defined^*, the phylogenetic tree *T^defined^*, and the site-rate heterogeneity model *ρ_i_* were subsequently used to simulate the alignment *D_i_* with *m_i_* sequences and sequence length *l_i_* using the AliSim program.

**Figure 1:**
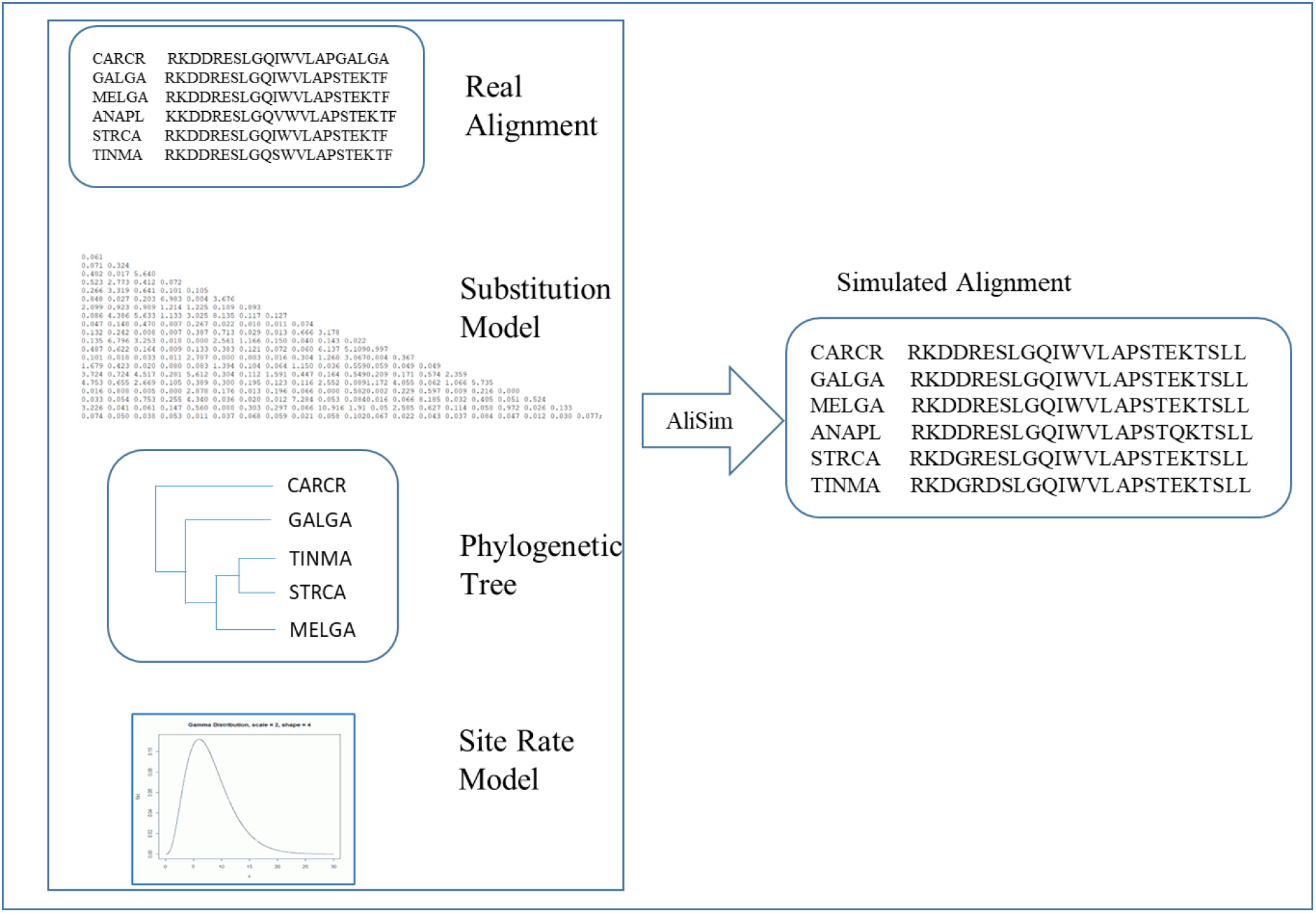
The procedure to simulate an alignment using parameters from a real alignment, a predefined amino acid substitution model, a predefined phylogenetic tree, and a predefined site rate model.

To generate a dataset **D** of *N* alignments, we conducted the simulation procedure *N* times with *N* different real alignments.

### Model analyses and comparison

First, we calculated the Pearson correlation between the true substitution matrix and the estimated matrix. We also analyzed the difference between amino acid exchangeability coefficients of the true model and that of the estimated model.

Let *Q^sim^* be the AA substitution model estimated from a simulated training dataset. More precisely, let *Q^simN^* be the AA substitution model estimated from a simulated training dataset of *N* alignments, e.g., *Q*^*sim*100^ be the AA substitution model estimated from a simulated training dataset of 100 alignments.

Second, we compared the quality of estimated model *Q^sim^* and the true model *Q^defined^* in building maximum likelihood trees on testing alignments. Technically, for each testing alignment *A_i_* if the ML tree 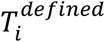 constructed with the true model *Q^defined^* is better (worse) than the tree 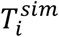 constructed with the estimated model *Q^sim^*, the true model *Q^defined^* is considered better (worse) than the estimated model *Q^sim^*. We also applied the significant test to examine if the tree 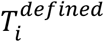 is significantly better (worse) than the tree 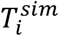. Technically, we performed the approximately unbiased (AU, *p* ≤ 0.05) test (Hidetoshi Shimodaira 2002) with 10.000 replicates using the CONSEL program (H. Shimodaira and Hasegawa 2002) to examine the significant likelihood difference between two trees.

Third, we compared the trees constructed with true models and that constructed with the estimated models using the Robinson-Foulds (RF) distance (Robinson and Foulds 1981). The RF distance is the number of clades which exists in one tree but not in the other tree. The normalized RF (nRF) distance is the RF distance normalized by dividing the RF distance to the number of clades, i.e., nRF ranges from 0 (i.e., two trees are identical) to 1 (i.e., two tree are completely different).

## Results

### Model analysis

We compared the true models Q.plant and NQ.plant with their corresponding simulated models Q^sim^.plant and NQ^sim^.plant estimated from different number of genes (see Figure 2). The models Q^sim^.plant and NQ^sim^.plant estimated from ≥ 100 genes have high correlations with the true models. For example, the correlation between the true model Q.plant (NQ.plant) and the estimated model Q^sim100^.plant (NQ^sim100^.plant) is 0.997 (0.999). We note that the correlations between the true models and models estimated from the genome datasets of 200 or 250 alignments are close to 1.

**Figure 2:**
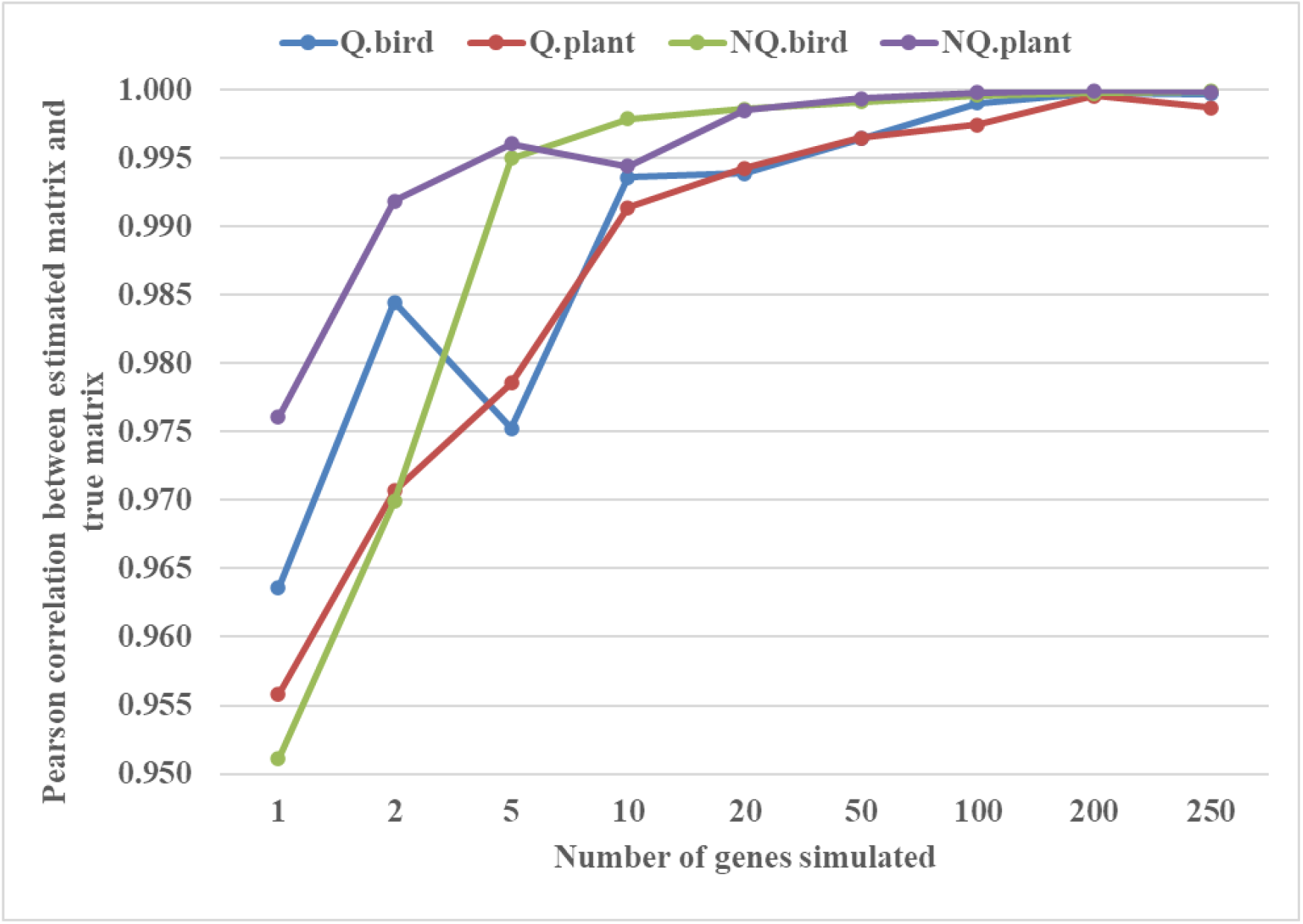
The Pearson correlations between the true substitution matrix and the estimated matrix for the bird and plant genome datasets with different number of genes. For genome datasets with ≥ 100 genes, the estimated models are very close to the true models.

We observed similar results for the bird datasets. The time reversible models Q^sim^.bird estimated with ≥ 100 genes have high correlations with the true model Q.bird (i.e., 0.999 for 100 genes; 0.9997 for 200 genes, and 0.9999 for 250 genes). The correlations between the true model and the estimated models with ≤ 10 genes are considerably low. For the time non-reversible models, the correlations between the true model NQ.bird and the estimated models NQ^sim^.bird with 100, 200, and 250 genes are 0.9991, 0.9995, 0.9997, 0.9998, respectively.

For more details, we counted the number of exchangeability coefficients in the estimated models that are at least two times or five times larger (smaller) than that in the true models. Table 1 shows the comparisons between the time reversible models Q^sim^.plant and Q.plant for the plant datasets; Q^sim^.bird and Q.bird for the bird datasets. The estimated models from datasets with less than 100 genes contain a considerable number of exchangeability coefficients that are at least two times difference in comparison with the true models. There are 14 (8) out of 189 elements in the simulated model Q^sim100^.plant that are at least two times greater (smaller) than that in the true model Q.plant. There are only few elements in the estimated models Q^sim200^.plant and Q^sim250^.plant that are two times or five times difference in comparison with that in the true model Q.plant. We observed similar results when comparing the estimated models Q^sim^.bird and the true model Q.bird. Figure 3 demonstrates the difference between estimated models Q^sim100^.plant and the true model Q.plant (3a), and between estimated models Q^sim100^.bird and the true model Q.bird (3b).

**Table 1:** The number of exchangeability coefficients that 2 times larger (2x), 5 times larger (5x), 2 times smaller (−2x), and 5 times smaller (−5x) when comparing time reversible models Q.plant with Q^sim100^.plant; and Q.bird with Q^sim100^.bird.

|  |  | Number of genes simulated |  |  |  |  |  |  |  |  |
| --- | --- | --- | --- | --- | --- | --- | --- | --- | --- | --- |
| Comparing models | Larger/Smaller | 1 | 2 | 5 | 10 | 20 | 50 | 100 | 200 | 250 |
| $Q^{\text{sim}}.\text{plant}$<br>vs.<br>$Q.\text{plant}$ | 2x | 85 | 62 | 63 | 35 | 30 | 13 | 14 | 10 | 6 |
|  | 5x | 75 | 51 | 42 | 17 | 18 | 5 | 2 | 1 | 4 |
|  | -2x | 24 | 27 | 18 | 21 | 15 | 12 | 8 | 6 | 3 |
|  | -5x | 12 | 9 | 6 | 6 | 3 | 3 | 2 | 6 | 2 |
| $Q^{\text{sim}}.\text{bird}$<br>vs.<br>$Q.\text{bird}$ | 2x | 81 | 63 | 67 | 50 | 38 | 28 | 16 | 12 | 9 |
|  | 5x | 76 | 56 | 54 | 37 | 22 | 12 | 5 | 2 | 1 |
|  | -2x | 27 | 19 | 13 | 11 | 5 | 2 | 0 | 1 | 1 |
|  | -5x | 8 | 2 | 2 | 4 | 0 | 0 | 0 | 0 | 0 |

**Figure 3:**
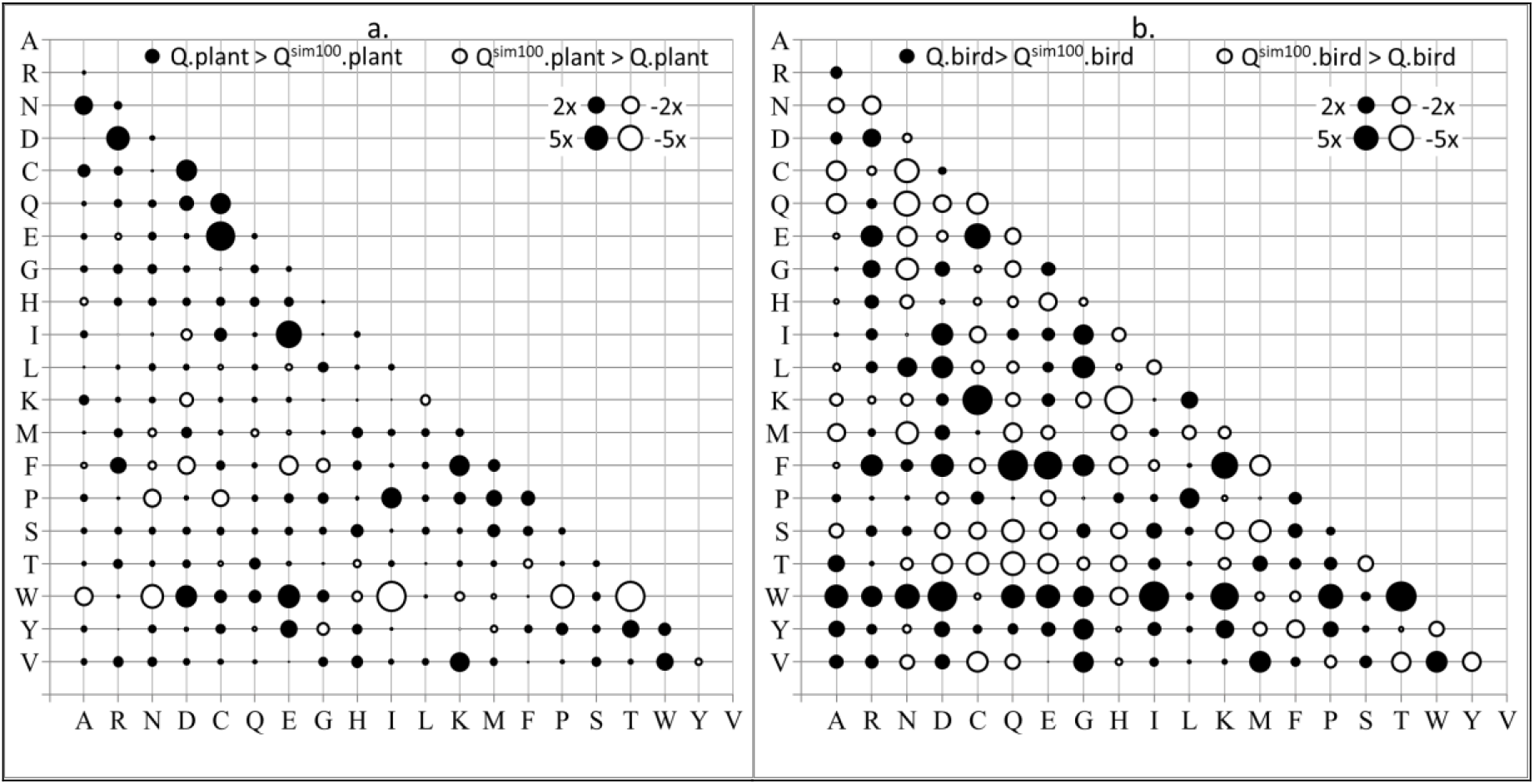
The bubble plots show relative differences between amino acid exchangeability coefficient in Q.plant and Q^sim100^.plant (a); and between Q.bird and Q^sim100^.bird (b). Both Q^sim100^.plant and Q^sim100^.bird were estimated from 100 simulated genes. Notations: 2x (5x) indicates that the exchangeability coefficient between two models is at least 2 times (5 times) difference.

We also compared the time non-reversible models NQ^sim^.plant and NQ^sim^.bird with their corresponding true models NQ.plant and NQ.bird (see Table 2). The NQ^sim100^.plant (NQ^sim200^.plant) model consists of 42 (23) exchangeability rates that are at least two times difference from that in the true model NQ.plant. Similarly, the NQ^sim100^.bird (NQ^sim200^.bird) model contains 36 (10) exchangeability rates that are at least two times difference from that in the true model NQ.bird. As the time non-reversible models consist of more free parameters than the time reversible models, more training data are required to estimate time non-reversible models so that they are close to the true models. Figure 4 demonstrates the difference between NQ^sim100^.plant and NQ.plant models (4a), and between NQ^sim100^.bird and NQ.bird models (4b).

**Table 2:** The number of exchangeability rates that 2 times larger (2x), 5 times larger (5x), 2 times smaller (−2x), and 5 times smaller (−5x) when comparing time non-reversible models NQ.plant with NQ^sim^.plant; and NQ.bird with NQ^sim^.bird..

|  |  | Number of genes simulated |  |  |  |  |  |  |  |  |
| --- | --- | --- | --- | --- | --- | --- | --- | --- | --- | --- |
| Comparing models | Larger/Smaller | 1 | 2 | 5 | 10 | 20 | 50 | 100 | 200 | 250 |
| NQ<sup>sim</sup>.plant<br>vs.<br>NQ.plant | 2x | 203 | 152 | 134 | 136 | 83 | 56 | 42 | 23 | 22 |
|  | 5x | 193 | 142 | 114 | 119 | 60 | 42 | 27 | 16 | 18 |
|  | -2x | 52 | 58 | 49 | 48 | 37 | 24 | 25 | 15 | 9 |
|  | -5x | 24 | 22 | 12 | 12 | 8 | 8 | 5 | 5 | 5 |
| NQ<sup>sim</sup>.bird<br>vs.<br>NQ.bird | 2x | 285 | 215 | 141 | 102 | 93 | 52 | 36 | 10 | 10 |
|  | 5x | 277 | 206 | 133 | 82 | 70 | 31 | 17 | 7 | 5 |
|  | -2x | 35 | 55 | 53 | 49 | 37 | 24 | 12 | 10 | 5 |
|  | -5x | 13 | 24 | 13 | 10 | 5 | 2 | 2 | 1 | 2 |

**Figure 4:**
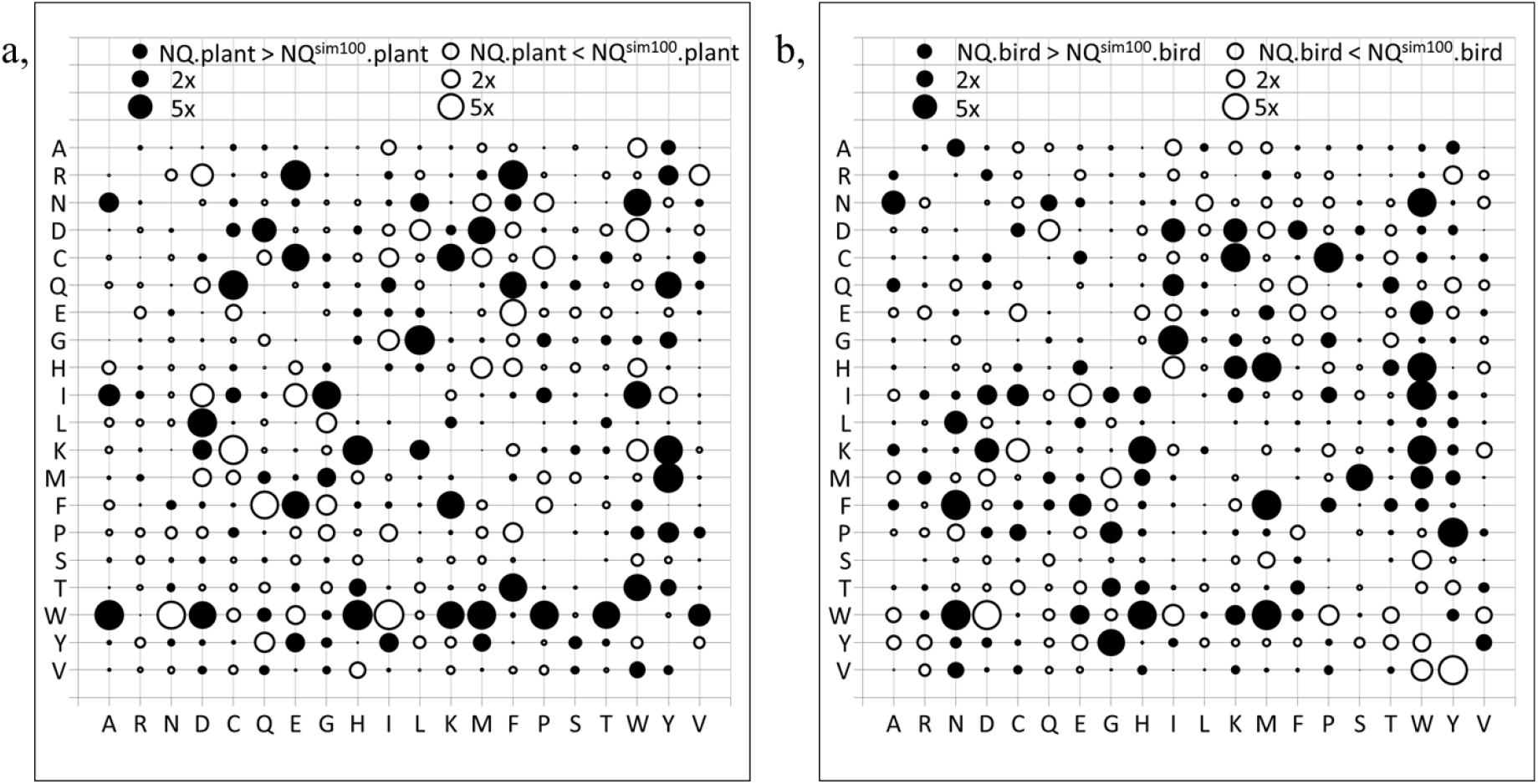
The bubble plots show relative differences between amino acid exchangeability rates in NQ.plant and NQ^sim100^.plant (a); and between NQ.bird and NQ^sim100^.bird (b). Both NQ^sim100^.plant and NQ^sim100^.bird were estimated from 100 simulated genes. Notations: 2x (5x) indicates that the exchangeability coefficient between two models is at least 2 times (5 times) difference.

### Likelihood comparison

We used the Bayesian information criterion (BIC) (Schwarz 2007) to examine the performance of estimated models in building maximum likelihood trees (see Figure *5*). We re-utilized the test datasets for plant and bird from the QMaker’s paper (Minh et al. 2021) which including 308 genes of plant and 500 genes of bird. For each of 308 genes of plant, the Q^sim^.plant (NQ^sim^.plant) model is better (worse) than the true model Q.plant (NQ.plant) if the ML tree constructed with Q^sim^.plant (NQ^sim^.plant) is better (worse) than that constructed with Q.plant (NQ.plant).

**Figure 5:**
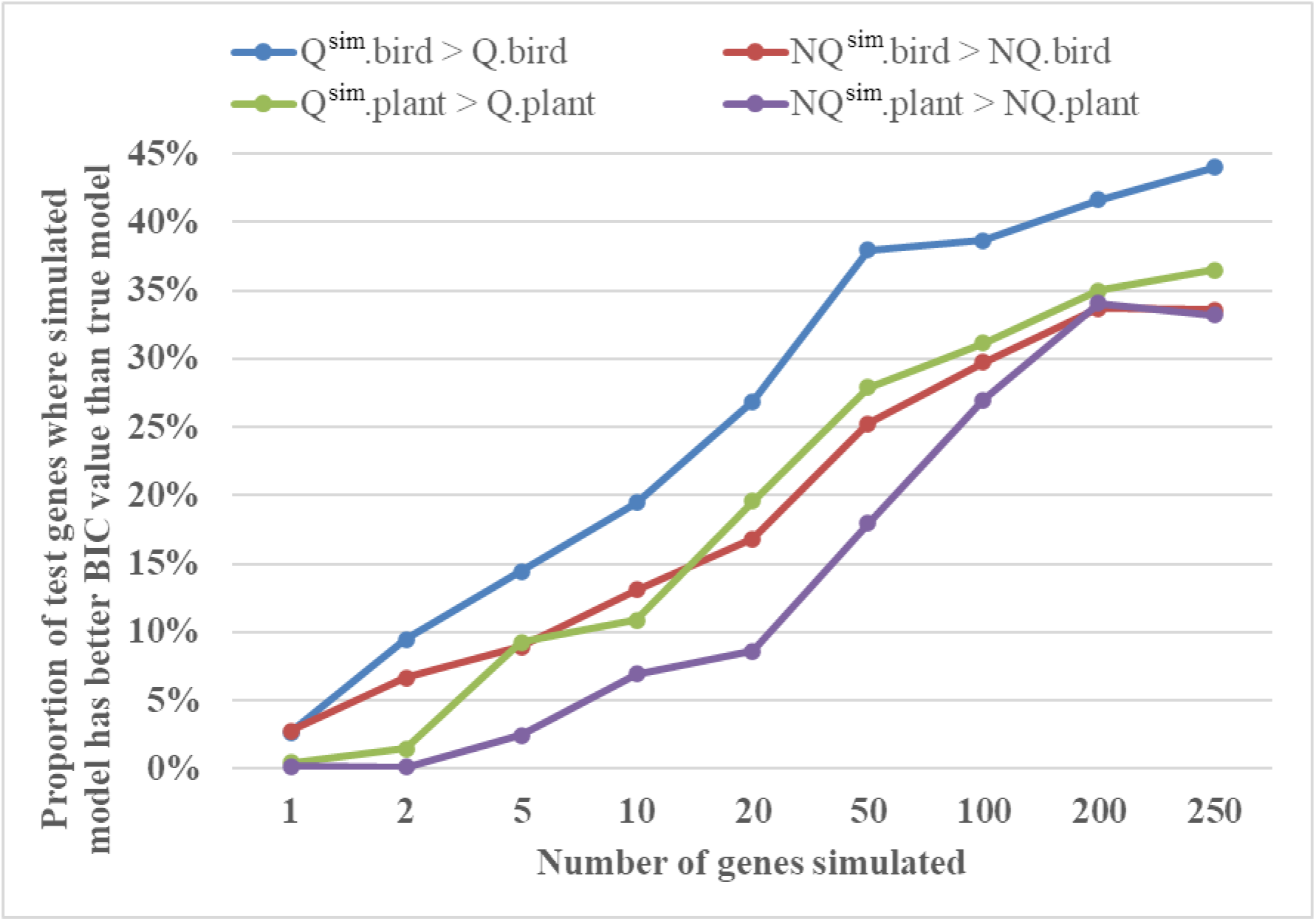
The performance of estimated and true models in building ML trees. The proportion of testing alignments where estimated model yields better BIC value than true model.

Figure *5* shows that the estimated models for plant with 10 or 20 genes were much worse than the true models in building ML trees. However, the plant models estimated with ≥ 100 genes performed considerably well in building ML trees in comparison with the true plant models. For example, the model Q^sim100^.plant (NQ^sim100^.plant) was better than the true model Q.plant (NQ.plant) in about 31.1% (27.01%) of testing alignments.

We obtained similar results for the bird datasets. The estimated model Q^sim100^.bird (Q^sim200^.bird) was better than the true model Q.bird in 38.6% (41.6%) of the testing alignments. The Q^sim^.bird models estimated from small datasets including 10 or 20 genes were much worse than the true model Q.bird. For the time non-reversible models, the estimated models NQ^sim100^.bird and NQ^sim200^.bird outperformed the true model NQ.bird in 31.4% and 33.2% of testing alignments, respectively.

We used the AU (*p* ≤ 0.05) test (Hidetoshi Shimodaira 2002) to determine if the likelihood difference between trees constructed with two different models is significant. Figure 6a shows the results for the plant datasets that the true models and the estimated models with ≥ 100 genes were not significantly better than each other for a majority of testing alignments. For example, Q.plant was significantly better than Q^sim100^.plant in 19.16% of the testing alignments. The estimated Q^sim100^.plant was significantly better than the true model Q.plant in 3.31% of the testing alignments. Thus, for 77.53% of the testing alignments, Q.plant and Q^sim100^.plant were not significantly better than each other.

**Figure 6:**
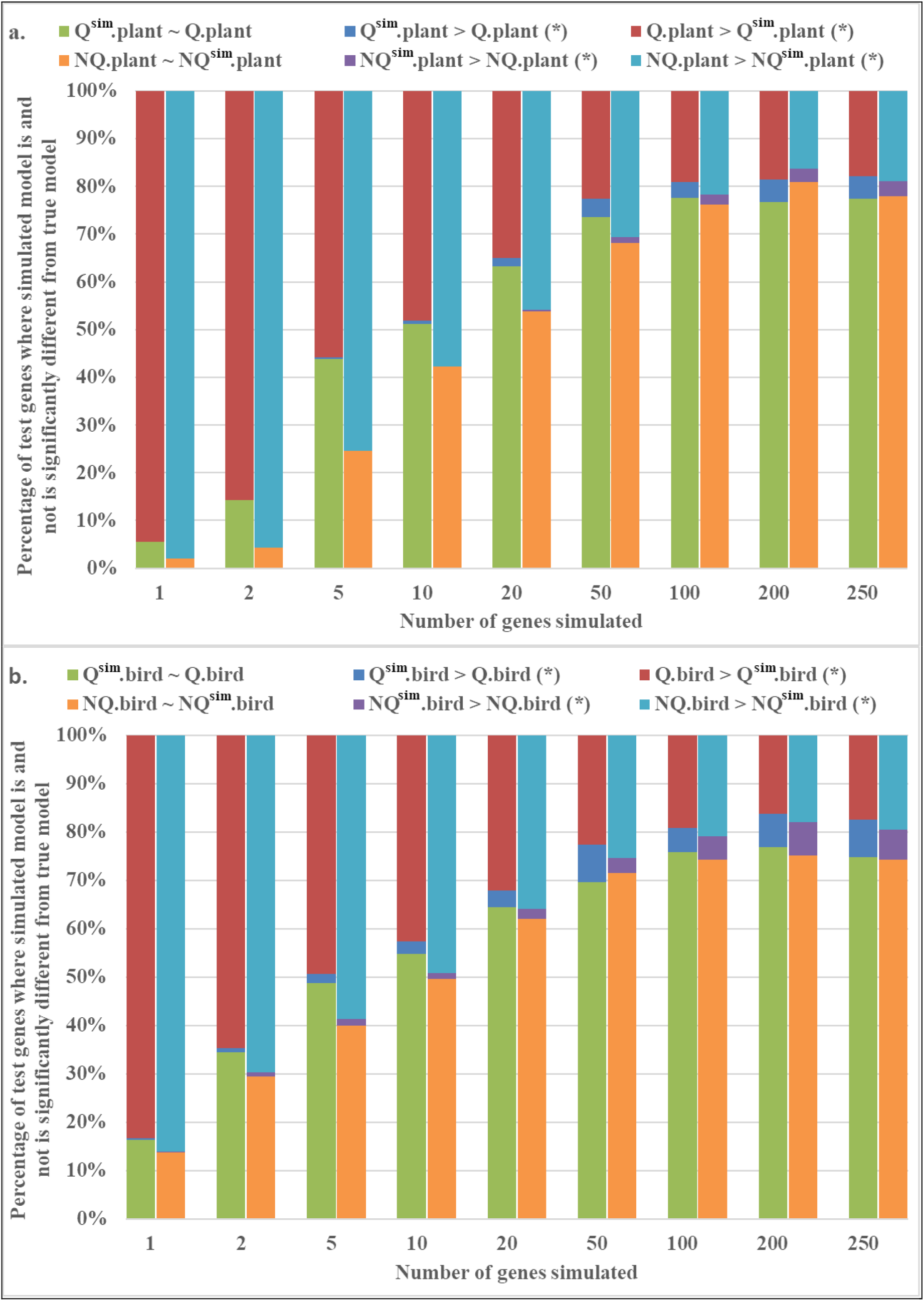
The AU tests (*p* ≤ 0.05) for the plant (a) and bird (b) dataset. Given a testing alignment, the AU test determines if the likelihoods of two trees constructed using the true and simulated models are significantly different. Q^sim^.plant ~ Q.plant: two trees constructed using two models Q^sim^.plant and Q.plant are not significantly different in term of likelihood. Q^sim^.plant > Q.plant (*): tree constructed using Q^sim^.plant has significantly better likelihood than tree constructed using Q.plant. Q.plant > Q^sim^.plant (*): tree constructed using Q.plant has significantly better likelihood than tree constructed using Q^sim^.plant. The same notion is applied to models of bird.

Similar results were observed when analyzing the bird datasets (Figure 6b). The AU test suggests that the true model Q.bird was only significantly better than the simulated model Q^sim100^.bird in 19.28% of the testing alignments. The simulated model Q^sim100^.bird was significant better than the true model Q.bird in 4.96% of the testing alignments. Thus, for 75.76% of the testing alignments, the Q.bird and Q^sim100^.bird were not significantly better than each other. Comparing the time non-reversible model NQ.bird and estimated models NQ^sim^.bird show that the true model NQ.bird and the estimated models with ≥ 100 genes were not significantly better than each other for more than 75% of testing alignments.

### Topology comparison

We examined the topological difference between trees reconstructed with the true models and that with the estimated models. The average nRF distances (see Table *3*) show that trees constructed with the true models and those with the estimated models are mostly identical. For plant alignments, trees inferred using the estimated models with ≥ 20 genes are identical with trees constructed with the true model. For the bird dataset, the average nRF distances between trees constructed with the true models and those inferred using the estimated models with ≥ 100 alignments are close to zero.

**Table 3:**
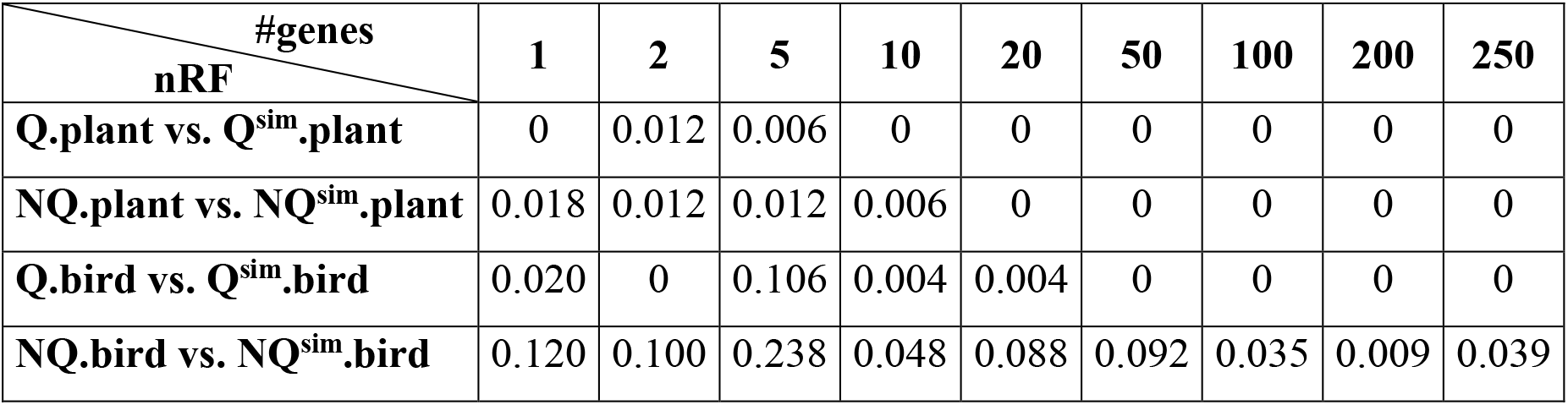
The average nRF distance between trees constructed with the true models and the estimated models. A *nRF* = 0 means that trees constructed with the true model and the estimated model are topologically identical.

| $\frac{\text{nRF}}{\text{\#genes}}$ | 1 | 2 | 5 | 10 | 20 | 50 | 100 | 200 | 250 |
| --- | --- | --- | --- | --- | --- | --- | --- | --- | --- |
| **Q.plant vs. Q<sup>sim</sup>.plant** | 0 | 0.012 | 0.006 | 0 | 0 | 0 | 0 | 0 | 0 |
| **NQ.plant vs. NQ<sup>sim</sup>.plant** | 0.018 | 0.012 | 0.012 | 0.006 | 0 | 0 | 0 | 0 | 0 |
| **Q.bird vs. Q<sup>sim</sup>.bird** | 0.020 | 0 | 0.106 | 0.004 | 0.004 | 0 | 0 | 0 | 0 |
| **NQ.bird vs. NQ<sup>sim</sup>.bird** | 0.120 | 0.100 | 0.238 | 0.048 | 0.088 | 0.092 | 0.035 | 0.009 | 0.039 |

## Discussion

The maximum likelihood methods have been proposed to estimate both time reversible and time non-reversible models from genome datasets. The true AA substitution models are unknown for real data; therefore, we are unable to evaluate the quality of estimated models in comparison with the true models based on the real data. In this paper, we compared the estimated models and the true models based on the simulated data. Our study based on the simulated genome datasets with different number of genes revealed that both time reversible and time non-reversible models estimated by the ML methods from large genome datasets (i.e., datasets with 100 or more genes) are closely related with the true models.

The estimated models from large datasets performed well in building maximum likelihood phylogenetic trees. The significant tests showed that the models derived from large genome datasets and the true models are not significantly better than each other for a majority of testing alignments. The tree topologies constructed with the estimated models and that inferred with the true models are nearly identical. As the true models are not available, the models estimated from large datasets can be reliably used in studying the evolution of protein sequences.

## Acknowledgements

This research was funded by the Vietnam National Foundation for Science and Technology Development (NAFOSTED; [102.01.2019.06 to CCD and LSV]).

## Author Contributions

CCD and LSV designed the study. TNH implemented the script and carried out the experiments. TNH drafted the manuscript. LSV and CCD revised the manuscript.

## Data availability

The datasets and scripts used in this paper are available at https://doi.org/10.6084/m9.figshare.22336384.

